# Long-term depression is independent of GluN2 subunit composition

**DOI:** 10.1101/270074

**Authors:** Jonathan M. Wong, John A. Gray

## Abstract

NMDA receptors (NMDARs) mediate major forms of both long-term potentiation (LTP) and long-term depression (LTD) and understanding how a single receptor can initiate both phenomena remains a major question in neuroscience. A prominent hypothesis implicates the NMDAR subunit composition, specifically GluN2A and GluN2B, in dictating the rules of synaptic plasticity. However, studies testing this hypotheses have yielded inconsistent and often contradictory results, especially for LTD. These inconsistent results may be due to challenges in the interpretation of subunit-selective pharmacology and in dissecting out the contributions of differential channel properties versus the interacting proteins unique to GluN2A or GluN2B. In this study, we address the pharmacological and biochemical challenges by utilizing a single-neuron genetic approach to delete NMDAR subunits in both male and female conditional knock-out mice. In addition, emerging evidence that non-ionotropic signaling through the NMDAR is sufficient for NMDAR-dependent LTD allowed the rigorous assessment of unique subunit contributions to NMDAR-dependent LTD while eliminating the variable of differential charge transfer. Here we find that neither the GluN2A nor the GluN2B subunit is strictly necessary for either non-ionotropic or ionotropic LTD.

## Introduction

NMDARs play prominent roles in bidirectional synaptic plasticity, mediating major forms of both long-term potentiation (LTP) and long-term depression (LTD) (Collingridge et al., 1983; Dudek and Bear, 1992). Most NMDARs are heterotetramers containing two obligatory GluN1 subunits and two GluN2 subunits, with GluN2A and GluN2B being the predominant subunits in the mammalian forebrain, including the hippocampus (Gray et al., 2011). Because the functional and regulatory properties of NMDARs are largely determined by their GluN2 subunit composition (Cull-Candy and Leszkiewicz, 2004), many studies have explored the hypothesis that different NMDAR subunits dictate the rules of synaptic plasticity (Shipton and Paulsen, 2014), though results have been inconsistent and often contradictory, especially for studies of long-term depression (LTD).

There are a number of potential reasons for the inconsistencies in LTD studies. First, interpretation of GluN2 subunit-selective pharmacology is problematic. GluN2 subunit-selective antagonists are limited by poor subunit selectivity (e.g. the GluN2A “selective antagonist” NVP-AAM077 is only 5-fold selective over GluN2B) (Neyton and Paoletti, 2006), incomplete blockade (e.g. ifenprodil only reduces currents from pure GluN2B-containing receptors about 80%) (Fischer et al., 1997; Hatton and Paoletti, 2005; Gray et al., 2011), and complex effects on glutamate affinity (e.g. ifenprodil increases glutamate affinity and prolongs NMDAR synaptic currents) (Kew et al., 1996; Gray et al., 2011; Tovar and Westbrook, 2012). Second, recent evidence has demonstrated that a high proportion of synaptic NMDARs are triheteromeric, containing GluN2A and GluN2B (Gray et al., 2011; Rauner and Kohr, 2011; Tovar et al., 2013). These triheteromeric receptors are only modestly responsive to GluN2-selective pharmacology (Hatton and Paoletti, 2005), further complicating the interpretation of these studies. Finally, conventional knock-out (KO) studies of GluN2 subunits have serious limitations as the GluN1 and GluN2B KO mice die perinatally (Forrest et al., 1994; Kutsuwada et al., 1996) and broad deletion of NMDARs results in altered network activity (Li et al., 1994; Iwasato et al., 2000).

Here we utilized a single-neuron genetic approach to isolate individual GluN2 subunits and assess their contributions to LTD. This approach avoids both the network-wide disruptions found in previous genetic manipulations as well as the difficult-to-interpret subunit specific pharmacology. Importantly however, even the interpretation of the effects of pure GluN2A or GluN2B receptor populations on synaptic plasticity can be problematic. Specifically, are effects of pure GluN2 subunit populations related to large differences in charge transfer (including Ca^2+^) or to critical associations with their divergent intracellular C-terminal tails? The inability to separate these variables further limits interpretations of NMDAR subunit-specific plasticity. Recently however, NMDAR-mediated LTD has been shown to occur in the absence of ion flux through the NMDAR (Nabavi et al., 2013; Stein et al., 2015; Carter and Jahr, 2016; but see Babiec et al., 2014), providing the opportunity to rigorously examine the GluN2 subunit-dependence of LTD while eliminating charge transfer as a variable. Surprisingly, we show no dependence of GluN2 subunit composition on either non-ionotropic or ionotropic NMDAR-dependent LTD.

## Materials and Methods

### Animals and postnatal viral injection

Animals were housed according to IACUC guidelines at the University of California Davis. *Grin2a*^fl/fl^ (Gray et al., 2011), *Grin2B*^fl/fl^ (Mishina and Sakimura, 2007; Akashi et al., 2009), and *Grin1*^fl/fl^ mice (Li et al., 1994; Adesnik et al., 2008) are all as previously described. Neonatal (P0-1) mice of both sexes were stereotaxically injected with high-titer rAAV1-Cre:GFP viral stock (~1-5×1012 vg/ml) with coordinates targeting CA1 of hippocampus as previously described (Gray et al., 2011). Transduced neurons were identified by nuclear GFP expression. Cre expression was generally limited to the hippocampus within a sparse population of CA1 pyramidal neurons.

### Electrophysiology

Mice were anesthetized in isoflurane and decapitated. Brains were rapidly removed and placed in ice-cold sucrose cutting buffer, containing (in mM) 210 sucrose, 25 NaHCO_3_, 2.5 KCl, 1.25 NaH_2_PO_4_, 7 glucose, 7 MgCl_2_, 0.5 CaCl_2_. Transverse 300μm hippocampal slices were cut on a Leica VT1200 vibratome (Buffalo Grove, IL) in ice-cold cutting buffer. Slices were recovered in 32°C artificial cerebrospinal fluid (ACSF) solution, containing (in mM) 119 NaCl, 26.2 NaHCO_3_, 11 glucose, 2.5 KCl, 1 NaH_2_PO_4_, 2.5 CaCl_2_ and 1.3 MgSO_4_, for 1 hour before recording. Slices were transferred to a submersion chamber on an upright Olympus microscope, perfused in room temperature normal ACSF containing picrotoxin (0.1 mM) and saturated with 95%O_2_/5%CO_2_. CA1 neurons were visualized by infrared differential interference contrast microscopy and GFP+ neurons were identified by epifluorescence microscopy. Cells were patched with 3-5MΩ borosilicate pipettes filled with intracellular solution, containing (in mM) 135 cesium methanesulfonate, 8 NaCl, 10 HEPES, 0.3 Na-GTP, 4 Mg-ATP, 0.3 EGTA, and 5 QX−314 (Sigma, St Louis, MO). Excitatory postsynaptic currents (EPSCs) were evoked by electrical stimulation of Schaffer collaterals with a bipolar electrode (MicroProbes, Gaithersburg, MD). AMPAR-EPSCs were measured at a holding potential of −70 mV, and NMDAR-EPSCs were measured at +40 mV in the presence of 10 μM NBQX. LTD was induced using a standard low-frequency stimulation protocol of 900 stimuli at 1 Hz (15 min) and holding the neuron at −40mV. Series resistance was monitored and not compensated, and cells were discarded if series resistance varied more than 25%. All recordings were obtained with a Multiclamp 700B amplifier (Molecular Devices, Sunnyvale, CA), filtered at 2 kHz, digitized at 10 Hz.. Analysis was performed with the Clampex software suite (Molecular Devices, Sunnyvale, CA).

### Experimental design and statistical analysis

All data represents the mean ± SEM of n = number of neurons or pairs or neurons. With the exception of the drug titrations, a minimum of three mice were used per group. All experimental groups include both males and females. Data were analyzed using Clampfit 10.4 (Axon instruments) and Prism 7 software (GraphPad). LTD experiments were analyzed by averaging the final 10 minutes of the recording and normalizing as a percent of the baseline AMPAR-EPSC amplitude. Paired amplitude and decay data were analyzed with a paired two-tailed *t* test and comparisons of LTD experiments were analyzed by unpaired two-tailed *t* test both with p<0.05 considered significant.

## Results

NMDAR glycine-site antagonists, which prevent channel opening, provide a key means to study non-ionotropic LTD. 7-chlorokynuernic acid (7CK) is a competitive NMDAR glycine-site antagonist that we and others have previously used to examine non-ionotropic LTD (Nabavi et al., 2013; Dore et al., 2015; Stein et al., 2015; Carter and Jahr, 2016). However, at concentrations needed for complete NMDAR block in acute brain slices (100 μM), 7CK also significantly inhibits AMPAR-EPSCs **(Figure 1**, purple, 74.9 ± 6.0%, n=4) making whole cell LTD recordings challenging. Thus, we have characterized the use of L689,560 (L689), a competitive glycine-site antagonist with higher potency and selectivity than 7CK (Leeson et al., 1992; Grimwood et al., 1995). A dose response of L689 on acute hippocampal slices found rapid, complete block of NMDAR-EPSCs by 10 μM L689 **(Figure 1A)**, a concentration that blocks only ~10% of AMPAR-EPSCs **(Figure 1B,C**; 10 μM L689, 10.7 ± 4.3%, n=4).

**Figure 1.**
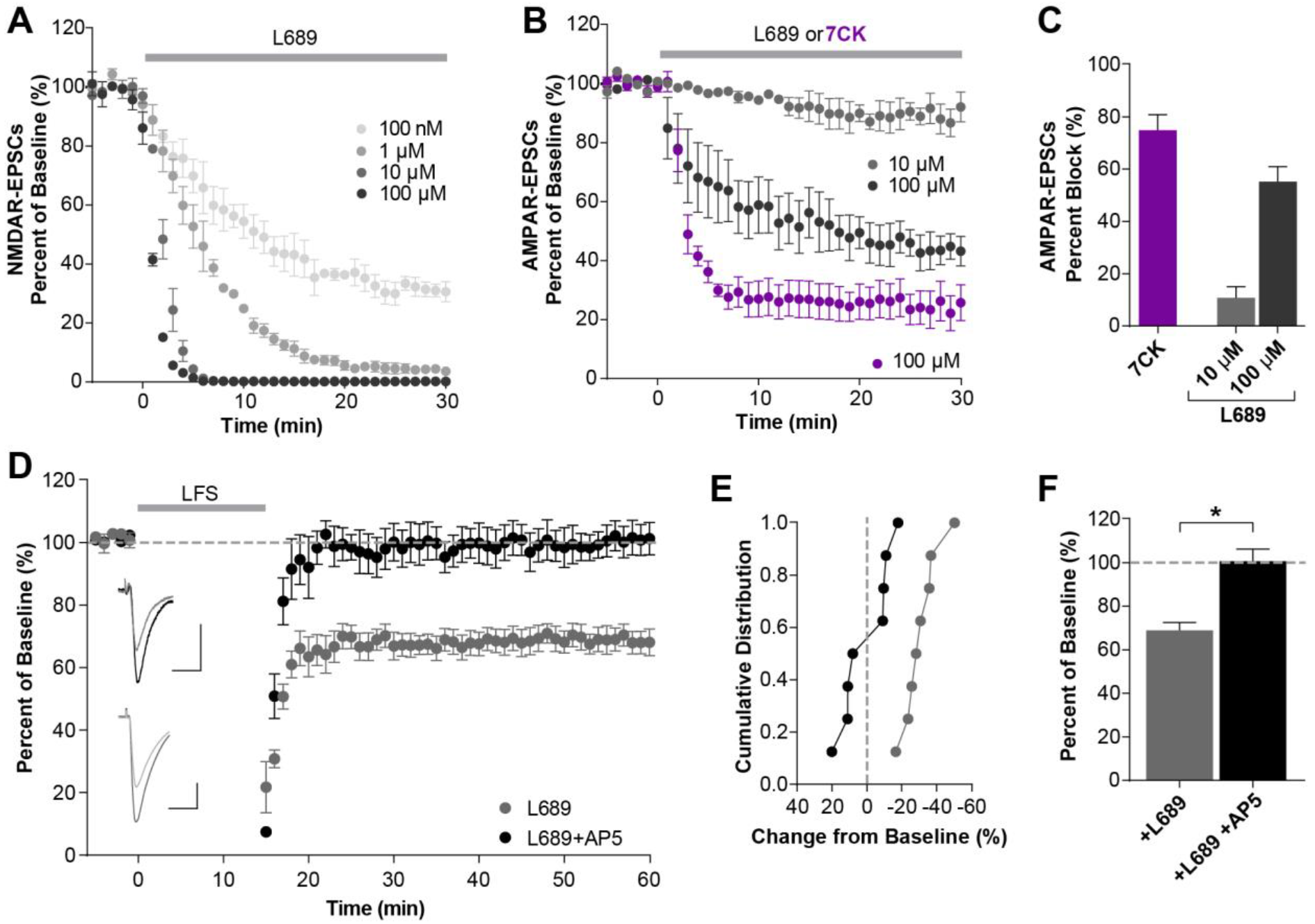
The NMDAR glycine-site antagonist L689 blocks NMDAR currents but not NMDAR-mediated LTD. **A**, Dose response of NMDAR-EPSC block by L689 in acute mouse hippocampal slices. NMDAR-EPSCs were fully inhibited by 10 μM and 100 μM L689 within 5 min (n=3 per dose). **B-C**, Inhibition of AMPAR-EPSCs by NMDAR glycine-site antagonists. (B) Time course of AMPAR-ESPC inhibition by 7CK and L689 normalized to baseline amplitude. (C) Percent block of AMPAR-EPSCs by 7CK and L689 averaging from 20-30 min after drug application. 100 μM 7CK and L689 inhibited AMPAR-EPSCs by 74.9 ± 6.0% and 55.2 ± 5.7%, respectively, while 10 μM L689 inhibited only 10.7 ± 4.3% (n=4 for each condition). **D-F**, Non-ionotropic NMDAR-mediated LTD occurs in the presence of 10 μM L689 and is blocked by 50 μM AP5. (D) Averaged whole cell LTD experiments and representative traces (10 ms, 50 pA). (E) Cumulative distribution of experiments in (D). (F) 10 μM L689 alone resulted in LTD (68.9 ± 3.6% of baseline, n=8). In contrast, addition of AP5 significantly inhibits this LTD (100.7 ± 5.5% of baseline, n=8) (*t*_(14)_=4.854, p=0.0003, *t* test). All data represents mean ± SEM.

### Non-ionotropic LTD is NMDAR-dependent

Consistent with 100 μM 7CK ((Nabavi et al., 2013; Stein et al., 2015), non-ionotropic LTD occurs in the presence of 10 μM L689 and remains NMDAR-dependent as it was blocked by concurrent incubation with the competitive glutamate-site antagonist AP5 **(Figure 1D-F**; L689, 68.9 ± 3.6%, n=8; +AP5, 100.7 ± 5.5%, n=8; *t*_(14)_=4.854, p=0.0003, *t* test). To further test the NMDAR-dependence of non-ionotropic LTD, we removed the obligatory GluN1 subunit in a sparse subset of CA1 pyramidal neurons by stereotaxic injection of adeno-associated virus, serotype 1 expressing a Cre recombinase GFP fusion protein (AAV1-Cre:GFP) into GluN1 conditional knockout mice (*Grin1*^fl/fl^) on postnatal day 0 (P0) **(Figure 2A)**. This mosaic deletion allows for simultaneous whole-cell recordings from Cre-expressing (Cre:GFP+) and untransfected neighboring cells, providing a rigorous comparison while controlling for presynaptic input. Consistent with our previous work (Gray et al., 2011), GluN1 deletion (ΔGluN1) results in a complete loss of NMDAR-EPSCs by P15 **(Figure 2B,C**; control, 82.1 ± 15.7 pA; ΔGluN1, 1.75 ± 0.53 pA; n=5, *t*_(4)_=5.021, p=0.007, paired *t* test). As expected, deletion of GluN1 prevented LTD in the presence of L689 **(Figure 2D-F**; control, 73.7 ± 3.5%, n=8; ΔGluN1, 99.8 ± 5.2%, n=8; *t*_(14)_=4.194, p=0.0009, *t* test). Together, these results demonstrate that non-ionotropic LTD is dependent on NMDARs.

**Figure 2.**
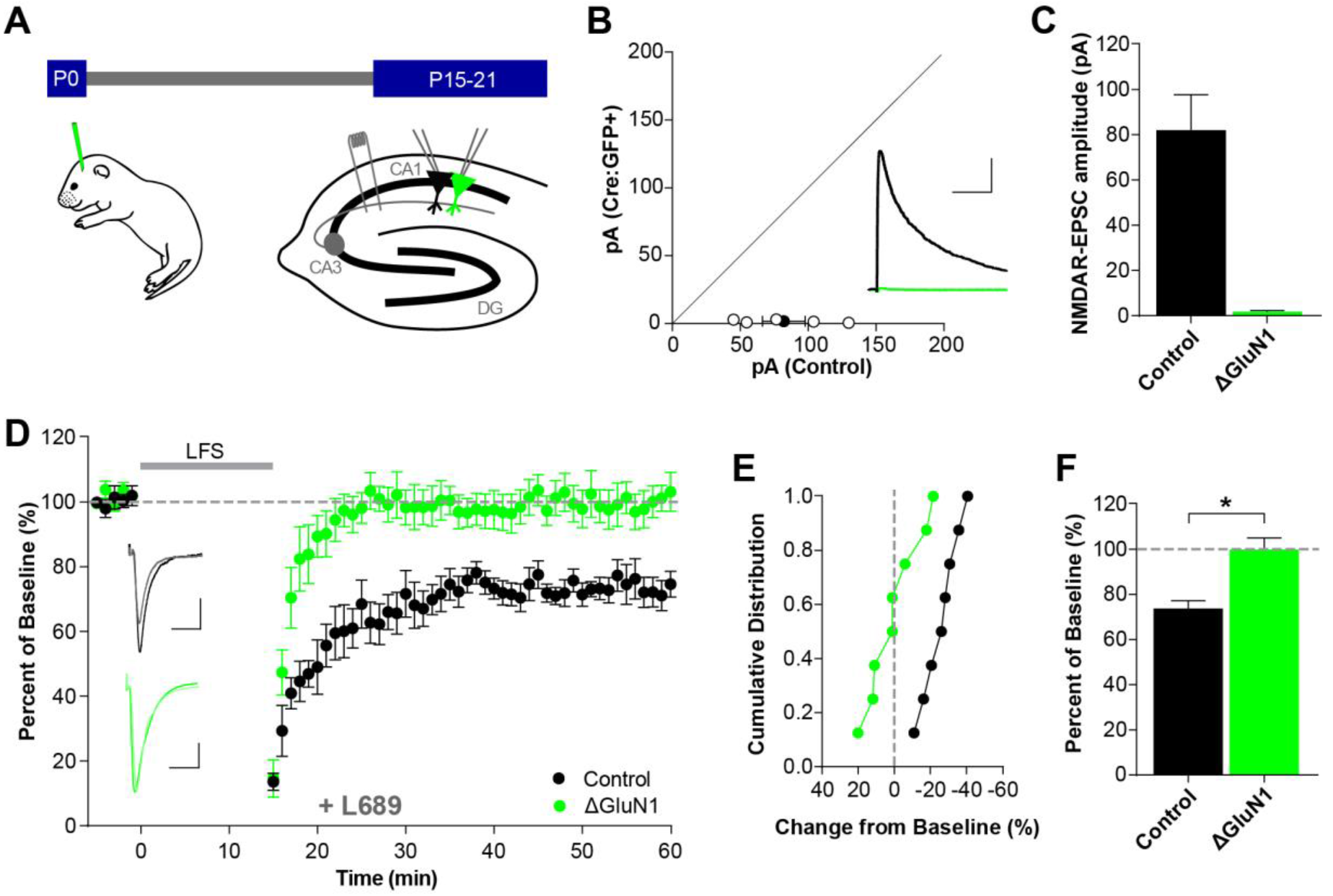
Single neuron deletion of GluN1 prevents non-ionotropic long term depression. **A**, Schematic of experimental preparation. Conditional knockout mice were injected with AAV1-Cre:GFP at P0. After 15-21 days, dual whole cell recordings were made from neighboring transduced and control neurons. **B-C**, NMDAR-EPSCs are eliminated by 15-21 days. (B) Scatterplot of individual neuron pairs (open circles) and averaged pair ± SEM (solid circle). Sample trace scale bars indicate (100 ms, 40 pA). (C) Average NMDAR-EPSC amplitudes for control (82.1 ± 15.7 pA, n=5) and Cre:GFP+ neurons (1.75 ± 0.53 pA, n=5); (*t*_(4)_=5.021, p=0.007, paired *t* test). **D-F**, Deletion of GluN1 prevents LTD. (D) Averaged whole cell LTD experiments and representative traces (10 ms, 50 pA). (E) Cumulative distribution of experiments in (D). (F) Average percent depression relative to baseline; control neurons (73.7 ± 3.5%, n=8), Cre:GFP+ neurons (ΔGluN1; 99.8 ± 5.2%, n=8), (*t*_(14)_=4.194, p=0.0009, *t* test).

### Non-ionotropic LTD is independent of GluN2 subtype

We next assessed the contribution of individual GluN2 subtypes to non-ionotropic LTD using single neuron deletion of GluN2A and GluN2B. As with GluN1, we performed simultaneous whole cell recordings of CA1 pyramidal neurons in *Grin2A*^fl/fl^ and *Grin2B*^fl/fl^ mice transduced with AAV1-Cre:GFP at P0. Deletion of GluN2A (ΔGluN2A) resulted in no change in the NMDAR-EPSC amplitude **(Figure 3A,B**; control, 102.8 ± 15.2 pA; ΔGluN2A, 96.4 ± 11.9 pA; n=6, *t*_(5)_=0.9913, p=0.367, paired *t* test) but a greatly prolonged EPSC decay **(Figure 3A,C**; control, 230.4 ± 8.5 ms; ΔGluN2A, 414.4 ± 13.3 ms; n=6, *t*_(5)_=13.35, p<0.0001, paired *t* test). This is consistent with our previous results (Gray et al., 2011) and represents a pure population of GluN2B-containing NMDARs. Deletion of GluN2A did not affect the expression of non-ionotropic LTD **(Figure 3D-F**; control, 77.1 ± 5.8%, n=6; ΔGluN2A, 65.1 ± 6.2%, n=6; *t*_(10)_=1.431, p=0.183, *t* test). Importantly, in interleaved experiments, AP5 continued to block LTD **(Figure 3G-I**; control, 97.0 ± 9.0%, n=6; ΔGluN2A, 96.7 ± 6.0%, n=6; *t*_(10)_=0.0274, p=0.979, *t* test) demonstrating that NMDAR-dependence is maintained.

**Figure 3.**
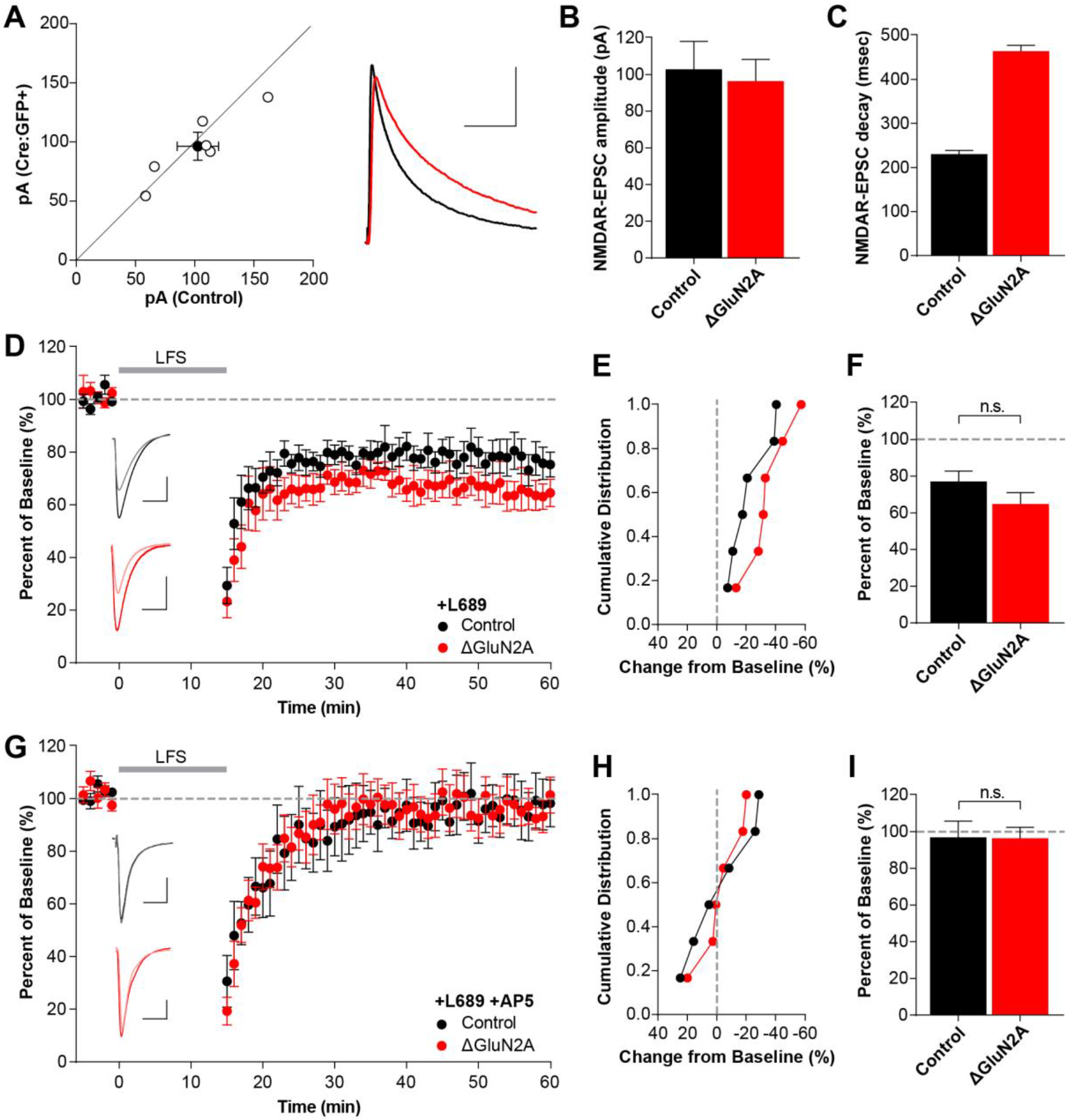
Single neuron deletion of GluN2A does not prevent non-ionotropic long term depression. **A-C**, Single neuron deletion of GluN2A. (A) Scatterplot of individual neuron pairs (open circles) and averaged pair ± SEM (solid circle). Sample trace scale bars indicate (100 ms, 40 pA). (B) Average NMDAR-EPSC amplitudes for control (102.8 ± 15.2 pA, n=6) and Cre:GFP+ neurons (96.4 ± 11.9 pA, n=6); p=0.48. (C) GluN2A deletion results in significantly longer decay kinetics (control 230.4 ± 8.5 ms, Cre:GFP+ 414.4 ± 13.3 ms; p<0.0001). **D-F**, GluN2A deletion does not block LTD. (D) Averaged whole cell LTD experiments and representative traces (10 ms, 50 pA). (E) Cumulative distribution of experiments in (D). (F) Average percent depression relative to baseline; control neurons (77.1 ± 5.8%, n=6), Cre:GFP+ neurons (ΔGluN2A; 65.1 ± 6.2%, n=6), (*t*_(10)_=1.431, p=0.183, *t* test) **G-I**, LTD after GluN2A deletion is still blocked by AP5. (G) Averaged whole cell LTD experiments and representative traces (10 ms, 50 pA). (H) Cumulative distribution of experiments in (G). (I) Summary graph of average percent depression relative to baseline; control neurons (97.0 ± 9.0%, n=6), Cre:GFP+ neurons (ΔGluN2A; 96.7 ± 6.0%, n=6), (*t*_(10)_=0.0274, p=0.979, *t* test).

Single neuron deletion of GluN2B (ΔGluN2B) resulted in a significant speeding of the NMDAR-EPSC decay time **(Figure 4A,C**; control, 233.7 ± 8.2 ms; ΔGluN2B, 79.0 ± 2.9 ms; n=6, *t*_(5)_=20.10, p<0.0001, paired *t* test) consistent with a pure population of GluN2A-containing NMDARs (Gray et al., 2011). Additionally, there was also a 30-40% reduction in the NMDAR-EPSC amplitude **(Figure 4A,B**; control, 90.1 ± 12.8 pA; ΔGluN2B, 58.1 ± 7.2 pA; n=6, *t*_(5)_=3.078, p=0.028, paired *t* test), as described previously (Gray et al., 2011). The simultaneous changes in NMDAR-EPSC amplitude and decay leads to a large decrease in charge transfer that could affect the interpretation of subunit dependence in LTD. However, deletion of GluN2B did not affect the expression of non-ionotropic LTD **(Figure 4D-F**; control, 76.1 ± 6.8%, n=8; ΔGluN2B, 74.3 ± 8.1%, n=9; *t*_(15)_=0.1662, p=0.870, *t* test) and this LTD remained NMDAR-dependent **(Figure 4G-I**; control 98.8 ± 7.3%, n=4; ΔGluN2B, 96.9 ± 8.6%, n=4; *t*_(6)_=0.1717, p=0.869, *t* test). Together, these results show that the expression of NMDAR-dependent non-ionotropic LTD is not dependent on the identity of the GluN2 subunit.

**Figure 4.**
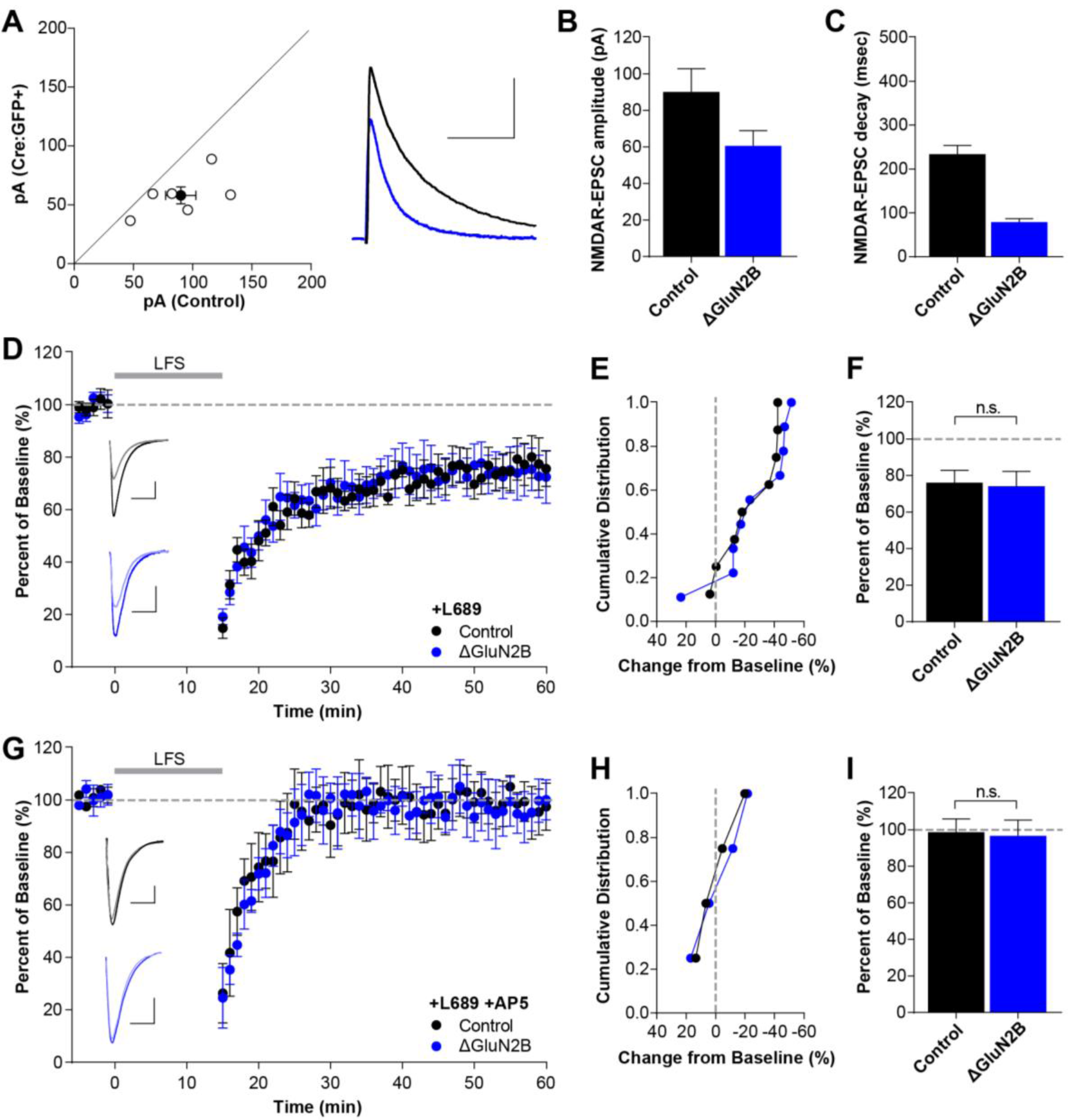
Single neuron deletion of GluN2B does not prevent non-ionotropic long term depression. **A-C**, Single neuron deletion of GluN2B. (A) Scatterplot of individual neuron pairs (open circles) and averaged pair ± SEM (solid circle). Sample trace scale bars indicate (100 ms, 40 pA). (B) Average NMDAR-EPSC amplitudes for control (90.1 ± 12.8 pA, n=6) and Cre:GFP+ neurons (58.1 ± 7.2 pA, n=6); p=0.016. (C) GluN2A deletion results in significantly faster decay kinetics (control 233.7 ± 8.2 ms, Cre:GFP+ 79.0 ± 2.9 ms; p<0.0001). **D-F**, GluN2B deletion does not block LTD. (D) Averaged whole cell LTD experiments and representative traces (10 ms, 50 pA). (E) Cumulative distribution of experiments in (D). (F) Average percent depression relative to baseline; control neurons (76.1 ± 6.8%, n=8), Cre:GFP+ neurons (ΔGluN2B; 74.3 ± 8.1%, n=9), (*t*_(15)_=0.1662, p=0.870, *t* test). **G-I**, LTD after GluN2B deletion is still blocked by AP5. (G) Averaged whole cell LTD experiments and representative traces (10 ms, 50 pA). (H) Cumulative distribution of experiments in (G). (I) Summary graph of average percent depression relative to baseline; control neurons (98.8 ± 7.3%, n=4), Cre:GFP+ neurons (ΔGluN2B; 96.9 ± 8.6%, n=4), (*t*_(6)_=0.1717, p=0.869, *t* test).

### Ionotropic LTD is independent of GluN2 subtype

The physiological relevance of non-ionotropic NMDAR-mediated LTD remains controversial (Gray et al., 2016). Thus, we examined the role of GluN2A and GluN2B is classical “ionotropic” LTD experiments in the absence of L689. Again, we found that both GluN2A-lacking and GluN2B-lacking neurons expressed LTD that was indistinguishable from control neurons **(Figure 5**; control, 75.5 ± 5.7%, n=8; ΔGluN2A, 68.9 ± 6.5%, n=9; ΔGluN2B, 81.6 ± 8.9%, n=9; control:ΔGluN2A, *t*_(15)_=0.7582, p=0.460, *t* test; control:ΔGluN2B, *t*_(15)_=0.5557, p=0.556, *t* test). Taken together, these findings provide rigorous evidence that NMDAR-mediated LTD is independent of GluN2 subunit composition.

**Figure 5.**
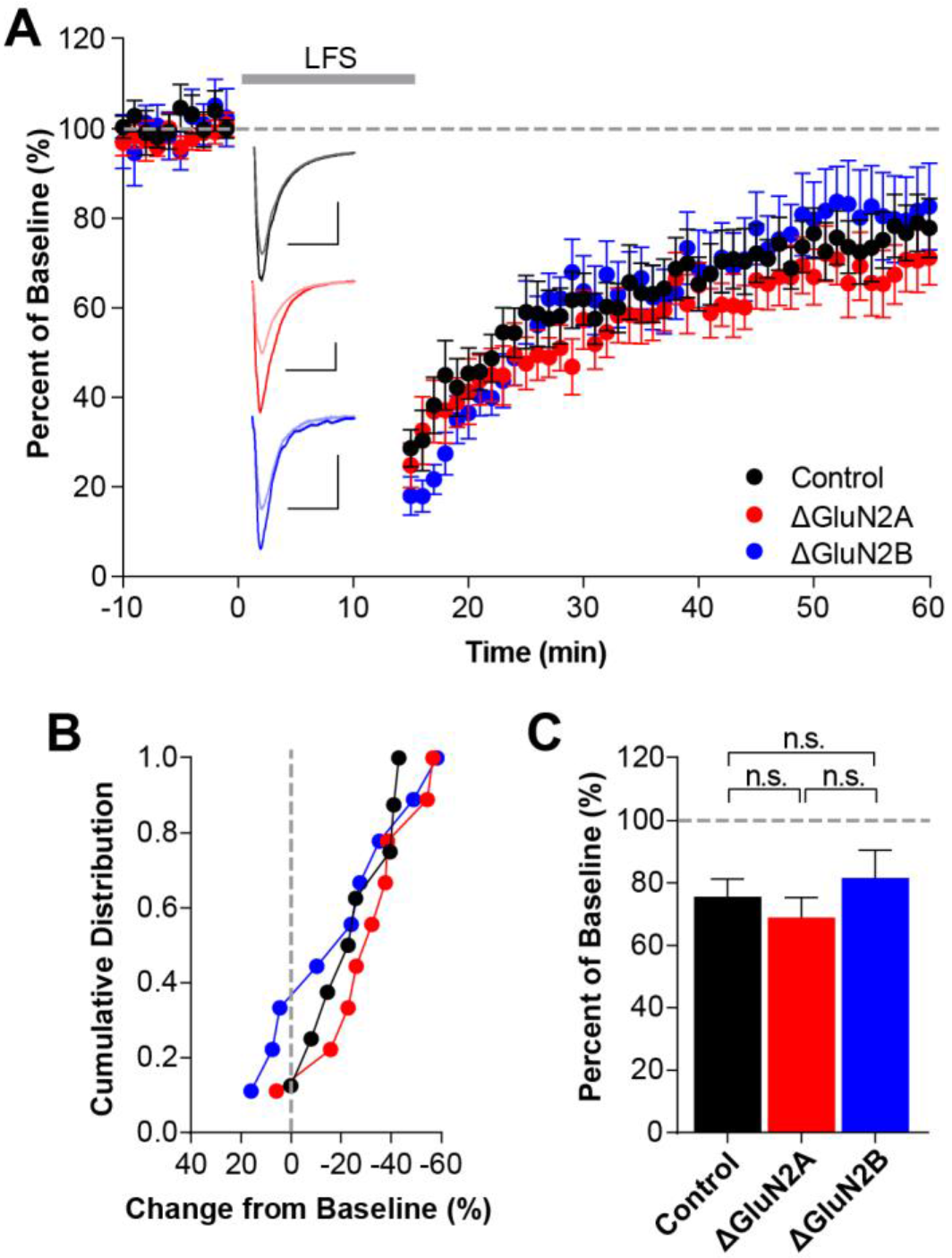
Single neuron deletion of either GluN2A or GluN2B does not prevent ionotropic long term depression. **A**, Averaged whole cell LTD experiments and representative traces (10 ms, 50 pA). **B**, Cumulative distribution of experiments in (A). **C**, Average percent depression relative to baseline; control neurons (75.5 ± 5.7%, n=8), ΔGluN2A neurons (68.9 ± 6.5%, n=9), ΔGluN2B neurons (81.6 ± 8.9%, n=9). There were no significant differences between control experiments and either GluN2A deletion (*t*_(15)_=0.7582, p=0.460, *t* test) or GluN2B deletion (*t*_(15)_=0.5557, p=0.556, *t* test), or between ΔGluN2A and ΔGluN2B (*t*_(16)_=1.151, p=0.267, *t* test).

## Discussion

Because major forms of both LTP and LTD are mediated by the NMDAR, it has long been hypothesized that the GluN2 subunit composition dictates the directionality of synaptic plasticity. This was an attractive hypothesis for a number of reasons. First, GluN2A and GluN2B confer distinct kinetic properties to synaptic NMDARs (Cull-Candy and Leszkiewicz, 2004) that could lead to the different levels of postsynaptic Ca^2+^ influx thought to underlie LTP and LTD (Dudek and Bear, 1992; Cummings et al., 1996; Yang et al., 1999; Rubin et al., 2005). Second, there is an activity-dependent developmental switch in synaptic NMDAR subunit composition in which predominantly GluN2B-containing NMDARs are replaced or supplemented by GluN2A (Sheng et al., 1994; Roberts and Ramoa, 1999). This subunit switch is thought to be a form of metaplasticity that alters the threshold and possibly the directionality of NMDAR-mediated synaptic plasticity (Quinlan et al., 1999; Dumas, 2005; Yashiro and Philpot, 2008; Gray et al., 2011). And third, GluN2A and GluN2B have long, highly divergent intracellular C-terminal domains that mediate an array of distinct protein-protein interactions that could be coupled to different downstream signaling pathways (Sanz-Clemente et al., 2013).

Numerous studies have set out to test the hypothesis that bidirectional plasticity is dictated by the GluN2 subunit composition, but their results have been inconsistent and conflicting, especially for LTD (reviewed by (Shipton and Paulsen, 2014). These inconsistent results are likely due to issues with GluN2 subunit-selective pharmacology (Neyton and Paoletti, 2006), thus we have utilized a mosaic genetic approach to delete NMDAR subunits in individual hippocampal neurons. Importantly however, genetically dissecting the relative roles of GluN2 subunits in synaptic plasticity is further complicated by altering two variables simultaneously: (1) differential postsynaptic Ca^2+^ dynamics between GluN2A and GluN2B, and (2) unique protein-protein interactions with their highly divergent C-terminal domains. The recent discovery of non-ionotropic NMDAR-mediated LTD (Nabavi et al., 2013), in which conformational changes in response to repetitive glutamate binding, but not channel opening or Ca^2+^ influx is posited to trigger LTD has provided a unique opportunity to reexamine the relative roles of GluN2A and GluN2B in synaptic plasticity. By removing Ca^2+^ influx as a variable, non-ionotropic LTD allows for a rigorous analysis of the subunit dependence of LTD. Our results here demonstrate conclusively that neither GluN2A nor GluN2B is strictly necessary for NMDAR-dependent LTD.

### Role of GluN2B in LTD

GluN2 subunit selective inhibition is confounded by poor selectivity, incomplete blockade, and complex effects on glutamate affinity (Kew et al., 1996). For GluN1/GluN2B receptors, ifenprodil and Ro 25-6881 antagonists are selective negative allosteric modulators that bind to the extracellular N-terminal domains (Hatton and Paoletti, 2005). Some studies have reported block of LTD by ifenprodil or Ro 25-6981 (Liu et al., 2004; Massey et al., 2004; Fox et al., 2006; Izumi et al., 2006; Gerkin et al., 2007; Ge et al., 2010; Dong et al., 2013; Izumi and Zorumski, 2015; Mizui et al., 2015; Yasuda and Mukai, 2015), though others report no effect (Hendricson et al., 2002; Bartlett et al., 2007; Li et al., 2007; Morishita et al., 2007; Kollen et al., 2008; Hanson et al., 2015; Yasuda and Mukai, 2015). However, these inhibitors display partial activity-dependence and only block a fraction (~80%) of synaptic GluN1/GluN2B diheteromers (Fischer et al., 1997; Hatton and Paoletti, 2005; Gray et al., 2011), which could result in variable effects based on drug concentration, slice activity, and pre-incubation time. Furthermore, N-terminal domain inhibitors only block about a quarter of the current in triheteromeric NMDARs (Hatton and Paoletti, 2005; Hansen et al., 2014) that make up a large proportion of synaptic NMDARs (Gray et al., 2011; Rauner and Kohr, 2011; Tovar et al., 2013). Another interesting consideration is that N-terminal domain inhibitors like ifenprodil decrease the glutamate dissociation rate (Kew et al., 1996; Gray et al., 2011; Tovar and Westbrook, 2012) that may have unknown effects on non-ionotropic LTD. For example, increasing glutamate affinity while preventing channel opening may promote non-ionotropic LTD, and one study reported that ifenprodil actually enhanced the magnitude of LTD (Hendricson et al., 2002). Taken together, the complexity of GluN2B-selective pharmacology makes firm conclusions on the role of GluN2B in LTD difficult.

### Role of GluN2A in LTD

For GluN2A-containing NMDARs, subunit-selective pharmacology is even more problematic. The most widely used antagonist, NVP-AAM007 (NVP), is a competitive glutamate-site antagonist that has only 10-fold selectivity for GluN2A over GluN2B (Neyton and Paoletti, 2006). As such, many LTD studies have used concentrations of NVP that antagonize a significant proportion of GluN2B (Liu et al., 2004; Massey et al., 2004; Izumi et al., 2006; Li et al., 2007). By titrating NVP to concentrations that block LTP, some groups found no inhibition of LTD (Liu et al., 2004; Gerkin et al., 2007; Ge et al., 2010), suggesting a key role for GluN2A in LTD, though other studies contradict this finding (Bartlett et al., 2007; Li et al., 2007). Given that NVP is a competitive glutamate site antagonist, NVP should consistently block LTD if only GluN2A is required; however, it remains unknown how NVP affects the triheteromeric receptors that predominate at earlier developmental points when LTD is most reliable. At “selective” concentrations, NVP should bind to the GluN2A glutamate site in triheteromers and block channel opening and LTP. However, it is unknown whether non-ionotropic LTD requires both glutamate sites to be occupied. Thus, continued glutamate binding to the GluN2B subunit in triheteromers could be sufficient to induce non-ionotropic LTD. Indeed, higher NVP concentrations consistently block LTD (Fox et al., 2006; Bartlett et al., 2007). Recently, more selective GluN2A inhibitors have been developed (e.g. TCN201) (Bettini et al., 2010; McKay et al., 2012) that block LTD (Izumi and Zorumski, 2015). Interestingly, these inhibitors have been shown to bind allosterically to the dimer interface between GluN1 and GluN2 (Hansen et al., 2012) which may impair conformational-based signaling. Overall, there remains no clear consensus on the role of GluN2A in LTD.

### Genetic studies of GluN2 subunits in LTD

In addition to pharmacological studies, a few genetic studies have addressed the GluN2 subunits in LTD. GluN2B KO mice die perinatally due to loss of suckling (Kutsuwada et al., 1996), but can survive by handfeeding. A loss of LTD was observed in hippocampal slices from three day old GluN2B KO mice (Kutsuwada et al., 1996), though at this age, a loss of GluN2B would result in a near complete loss of synaptic NMDARs (Gray et al., 2011). Selective deletion of GluN2B impaired LTD (Brigman et al., 2010) in 14-22 week old mice, though LTD required block of glutamate transporters to induce spillover, presumably to activate extrasynaptic receptors. Importantly, these studies were at the developmental time points that widely deviate from the standard LTD literature making generalization difficult. Interestingly, acute disruption of the interaction of GluN2B with PSD95 using a cell-permeable peptide reduced synaptic GluN2B levels and impaired LTP but had no effect on LTD (Gardoni et al., 2009), consistent with our findings that GluN2B is not necessary. Fewer studies have examined GluN2A, though germline GluN2A KO mice have normal NMDAR-dependent LTD in CA1 (Longordo et al., 2009; Kannangara et al., 2015).

### Mechanism of non-ionotropic LTD

The widely-accepted model for bidirectional synaptic plasticity mediated by NMDAR activation posits that large, rapid increases in synaptic Ca^2+^ leads to LTP and prolonged, modest increases in Ca^2+^ leads to LTD (Lisman, 1989; Malenka, 1994; Neveu and Zucker, 1996). This model has recently been challenged with the finding that repetitive glutamate binding to the NMDAR is sufficient to induce LTD and spine shrinkage, independent of Ca^2+^ influx (Nabavi et al., 2013; Stein et al., 2015; Carter and Jahr, 2016; Gray et al., 2016), though this remains controversial (Babiec et al., 2014). Importantly, a role for Ca^2+^ in the expression of LTD remains, as intracellular Ca^2+^ chelators inhibit non-ionotropic LTD (Nabavi et al., 2013). However, clamping intracellular Ca^2+^ at baseline concentrations while preventing Ca^2+^ elevations rescued the expression of non-ionotropic LTD (Nabavi et al., 2013). These findings suggest that non-ionotropic LTD involves glutamate-mediated conformational changes in the NMDAR (Dore et al., 2015).

Conformation-based signaling by the NMDAR suggests modulation of receptor interacting partner(s), and the long intracellular C-terminal tails of the GluN2 subunits were the most likely candidates. For example, the death-associated protein kinase 1 (DAPK1) competes with the binding of CaMKII to GluN2B promoting LTD over LTP (Goodell et al., 2017). However, our current results suggest that these interactions are no strictly necessary for LTD and that the minimum sufficient LTD signal is not based on the divergence of the GluN2 subunits. So, without Ca^2+^ influx or unique GluN2 interacting proteins, what could be the crucial receptor-proximal factor for LTD? Possibilities include shared interactions between GluN2A and GluN2B, interactions with GluN1, or transmembrane or extracellular interactions. For example, protein phosphatase 1 (PP1) is a key intermediary protein which is displaced from GluN1 following NMDA binding suggesting a GluN1-proximal mechanism (Dore et al., 2015). Further studies are needed to identify the minimum NMDAR determinates necessary for LTD and to examine whether ionotropic and non-ionotropic LTD are identical or parallel processes.

## Acknowledgements

This work was supported by NIH Grant K08MH100562 and a pilot grant through the UC Davis Alzheimer’s Disease Center P30AG010129. We thank Zaiyang “Sunny” Zhang and Haley Martin for their assistance with mouse breeding and genotyping.

